# Fully Desktop Fabricated Flexible Graphene Electrocorticography (ECoG) Arrays

**DOI:** 10.1101/2022.07.25.501414

**Authors:** Jia Hu, Ridwan Fayaz Hossain, Zahra S. Navabi, Alana Tillery, Michael Laroque, Preston D. Donaldson, Sarah L. Swisher, Suhasa B. Kodandaramaiah

## Abstract

Flexible Electrocorticography (ECoG) electrode arrays that conform to the cortical surface and record surface field potentials from multiple brain regions provide unique insights into how computations occurring in distributed brain regions mediate behavior. Current flexible ECoG devices require highly specialized microfabrication methods, precluding the ability to fabricate customizable and low-cost flexible ECoG devices easily. Here we present a fully desktop fabricated flexible graphene ECoG array. First, we synthesized a stable, conductive ink via liquid exfoliation of Graphene in Cyrene. Next, we have established a stencil-printing process for patterning the graphene ink via laser-cut stencils on flexible polyimide substrates. Benchtop tests indicate that the graphene electrodes have good conductivity of ∼ 1.1 × 10^3^ S·cm^-1^, flexibility to maintain their electrical connection under static bending, and electrochemical stability in a 15-day accelerated corrosion test. Chronically implanted graphene ECoG devices remain fully functional for up to 180 days, with average *in vivo* impedances of 24.72 ± 95.23 k Ω at 1 kHz. The ECoG device can measure spontaneous surface field potentials from mice under awake and anesthetized states and sensory stimulus-evoked responses. The stencil-printing fabrication process can be used to create Graphene ECoG devices with customized electrode layouts within 24 hours using commonly available laboratory equipment.

## INTRODUCTION

Simultaneous neural computations occurring in many brain regions and interactions between these regions mediate behavior [1]–[3]. Disruptions to these interactions are an indicator of neurological disorders such as Parkinson’s disease [4] and traumatic brain injury [5]. Minimally invasive electrocorticography (ECoG) electrode arrays have been used to measure surface field potential for recording neural activities from several distributed populations of neurons at the brain surface [6]. Functionally, ECoG electrode arrays covering large brain regions need to be flexible to make adequate contact with the complex, convex surface of the brain [6], [7]. To this end, flexible, silicon substrate-based [8] and polymer substrate-based ECoG devices [7]–[19] have been developed. However, these devices were made using highly specialized micro- or nanofabrication techniques which require training and specialized fabrication equipment. Further, most neuroscience laboratories require rapid and flexible design alterations to adapt to various experimental contexts, which is hard to achieve in traditional microfabrication procedures. To simplify the fabrication procedure, inkjet printing conductive materials such as silver nanoparticle inks [9] and conductive polymers like PEDOT:PSS [10] has been used to create flexible and reconfigurable ECoG electrode arrays. Currently, these approaches require expensive printers and still rely on specialized or microfabrication techniques to insulate the electrode [11].

This work introduces a fully-desktop fabricated stencil-printed, flexible graphene ECoG electrode array fabricated using commonly available tools in neuroscience laboratories. Graphene has become an attractive material for realizing neural interfaces [12]–[16] due to its high conductivity [17], flexibility [18], transparency [19], stability [20], and biocompatibility [18]. We first formulated a stable conductive ink based on exfoliating Graphene in Cyrene. The ink was stencil-printed on a flexible polyimide substrate via laser-cut stencils. A similar procedure was applied to pattern a sacrificial Pluronic layer, a tri-block copolymer for defining the exposed electrode pads. Subsequently, a silicone elastomer insulation layer was coated on the electrodes using a custom-built, microprocessor-controlled spin-coater to create functional ECoG electrode arrays. Most importantly, no aspect of the device fabrication needed access to specialized cleanroom facilities. The ECoG arrays exhibited excellent benchtop performance characteristics and *in vivo* recording capabilities. Using this methodology, the proposed devices can be fabricated within 24 hours and require a level of complexity and skill comparable to the assembly of tetrode devices [21]. The flexible Graphene ECoG devices were chronically implanted in mice and demonstrated the ability to record surface field potentials for up to 180 days.

## RESULTS

### Desktop fabricated flexible graphene ECoG electrodes

We sought to engineer a flexible graphene ECoG device that could be entirely fabricated using available tools and equipment in most neuroscience laboratories. Previous work has been done on stencil-printed graphene electronics [22]–[25], but fully desktop fabricated µ-ECoG arrays implanted in mice have not been developed. The fabrication procedure required first a graphene ink which can be used to build electrodes and retain electrical performance after chronic implantation. Second, we sought to establish a simple process for achieving reconfigurable electrode arrays for use in small animals. In this study, we used graphene exfoliated in Cyrene solvent to create a high-performance conductive ink that could be patterned on a polyimide film using a stencil-printing technique (**Fig. 1a**).

**Figure 1.**
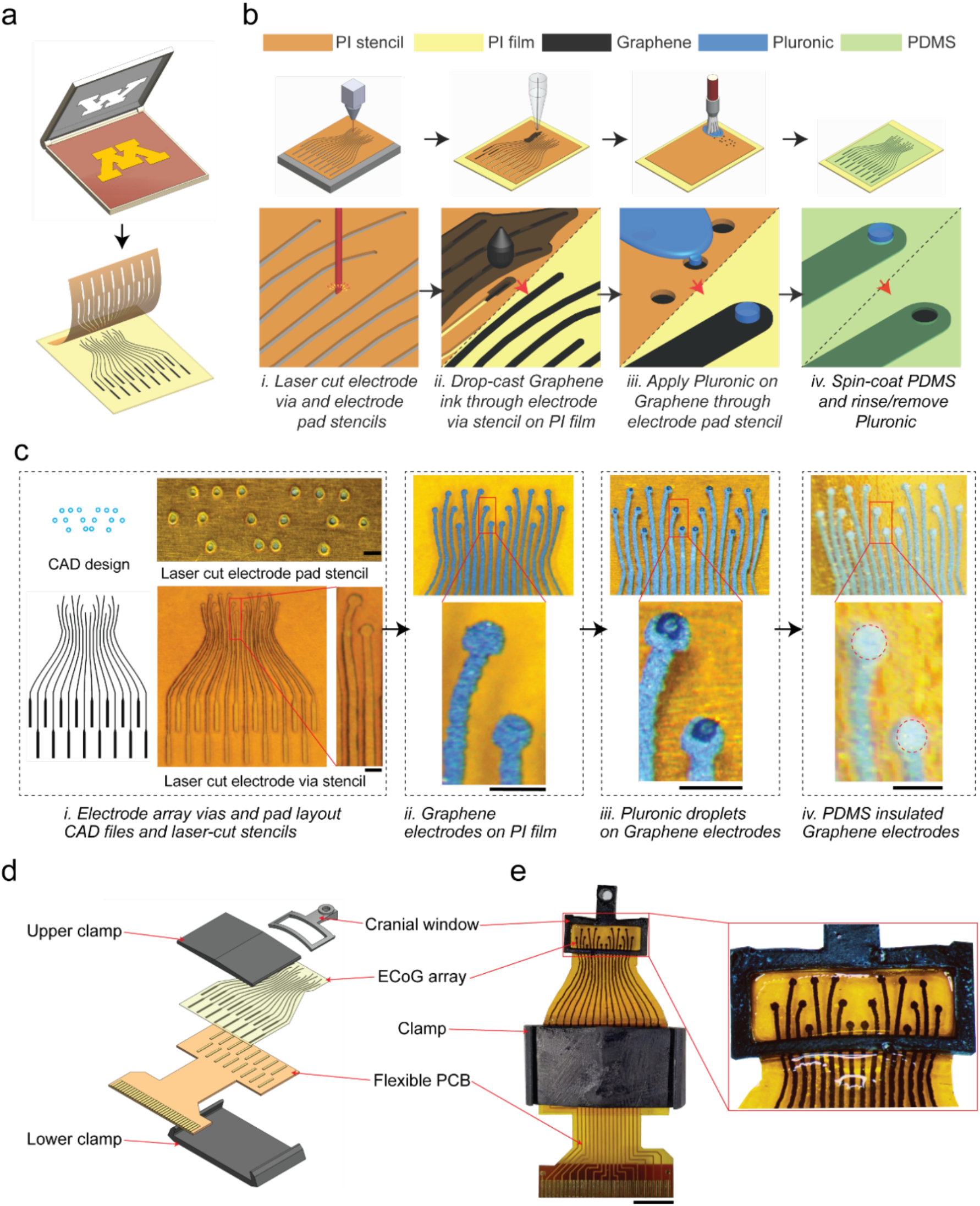
Stencil-printed flexible graphene ECoG array: **(a)** stencil-printing adapted from screen-printing to realize flexible neural interfaces. **(b)** Schematic of the graphene ECoG electrode array fabrication procedure. (i) Stencils are cut from insulating polyimide tape (PI) using a desktop laser; (ii) graphene ink was drop-cast via stencil on the PI film to pattern the electrode traces, followed by annealing; (iii) Pluronic is deposited through the electrode pad stencil on the graphene electrodes; (iv) Diluted PDMS is spin coated to form the insulating layer. After PDMS cures, Pluronic is rinsed away to expose the electrode pads. **(c)** (i) Computer-aided design (CAD) drawing and laser-cut stencils of electrode array vias and electrode pads. (ii) The patterned graphene electrode on PI film. (iii) Pluronic droplets on graphene electrodes. (iv) PDMS insulated graphene electrodes. Scale bars indicate 500 µm. **(d)** CAD rendering of the whole implant assembly. **(e)** Photograph of a fully assembled 16-channel graphene ECoG device. Scale bar indicates 5 mm.

The overall fabrication procedure is shown in **Fig. 1a**. In the first step, stencils for patterning the graphene electrodes and a sacrificial layer of Pluronic defining the exposed electrode pads were created using a desktop laser cutter (**Fig. 1a. i**). The desktop laser cutter achieved a minimum inter-electrode (center to center) distance of ∼ 400 µm. To reliably create isolated electrodes without any spurious interconnects, we used the inter-electrode distance of 500 µm in the stable design (**Fig. 1b, c**). This inter-electrode distance depended on the laser power and cutting speed of the laser cutter, which was experimentally optimized (**S.Fig. 1**).

The laser-cut stencil was overlaid on a flexible polyimide (PI) film, followed by drop-casting graphene ink to pattern the electrode arrays (**Fig. 1a. ii**). After drop-casting, the PI film with the stencil was annealed at a low temperature of ∼ 100 °C to evaporate the excess Cyrene solvent. Then, the stencil was stripped, and the electrode array was annealed at a high temperature of ∼ 300 °C (**Fig. 1c**). At room temperature, the second stencil for patterning the sacrificial Pluronic layer was overlaid on the substrate (**Fig. 1a. iii**). A fine paintbrush was used to apply Pluronic on the electrode pad areas (**Fig. 1a. iii**). After removing the stencil, the Pluronic was allowed to solidify overnight (**Fig. 1c. iii**), then the array was spin-coated with diluted silicone elastomer to create the insulating layer. Once the elastomer was cured, the electrode array was gently rinsed in warm water to remove the sacrificial Pluronic covering the electrode pads, resulting in an insulated electrode array with exposed electrode sites for interfacing with a printed circuit board (PCB) (**S. Fig. 1a**) and exposed electrode pads with an average diameter of ∼ 300 µm for interfacing the brain. Once the electrode arrays were fabricated, they were integrated into a 3D printed frame adapted from our previous work [9], [26], [27]. The PCB interface was bonded to the ECoG array using conductive epoxy (**S. Fig. 2a**) and mechanically reinforced using a custom 3D printed clamp (**Fig. 1d, e**). The fabrication procedure relied entirely on common equipment such as a laser cutter and stereolithographic 3D printers, ubiquitous in nearly every university maker space. The electrode annealing was performed using a laboratory hotplate. Spin-coating was performed using a homebuilt microprocessor-controlled spinner (**S. Fig. 5**), but off-the-shelf low-cost speed-controllable spin coaters under $1000 can also serve the purpose.

### Formulation of graphene inks and material characterization

We formulated three graphene inks for stencil fabrication by exfoliating graphene in Cyrene, Dimethylformamide (DMF) [28]–[31], and Cyclohexanone/Terpineol (C/T) [18], [32], [33] solvents, using a top-down liquid-phase exfoliation (LPE) technique through ultra-sonication (see **Methods** for details). graphene inks in DMF and C/T have already been used in optoelectronic, photovoltaic, and biomedical applications [18], [29]–[34]. More recently, Cyrene demonstrated near-ideal physical properties for graphite exfoliation and the production of graphene dispersions [35]. Graphene exfoliated in Cyrene for the same sonication time of 2 hours resulted in drop-cast films with sub-micrometer surface flakes, while graphene in DMF and C/T ink produced rougher films with much larger surface clusters (**Fig. 2a**). As shown in **S.Figure 6**, the Raman spectroscopy was performed on spin-cast films from the three ink samples. The I_d_/I_g_ ratio for highly disordered graphitic films like those generated from annealed graphene flakes increases with crystallite (i.e., flake) size [36]. The I_d_/I_g_ ratio for the Cyrene-exfoliated graphene sample is the lowest, indicating it has the least average distance between defects (< 1 nm), implying that this sample is composed of the smallest flakes of the three inks, which have been annealed into a mainly sp^2^ amorphous carbon structure [37]. Flake size is a determinant of the overall packing density, with a smaller flake size enhancing the inter-platelet connectivity, resulting in higher conductivity [18]. Further, smaller flake sizes are desirable for uniform patterning through stencils or inkjet printing if needed [18], [22]. The thickness of the drop-cast graphene inks were measured with a surface profiler, where the Cyrene+Gr, C/T+EC+Gr, DMF+EC+Gr films’ thickness were measured to be 5.84 µm, 12.93 µm, and 8.08 µm, respectively (**S. Fig. 7**).

**Figure 2.**
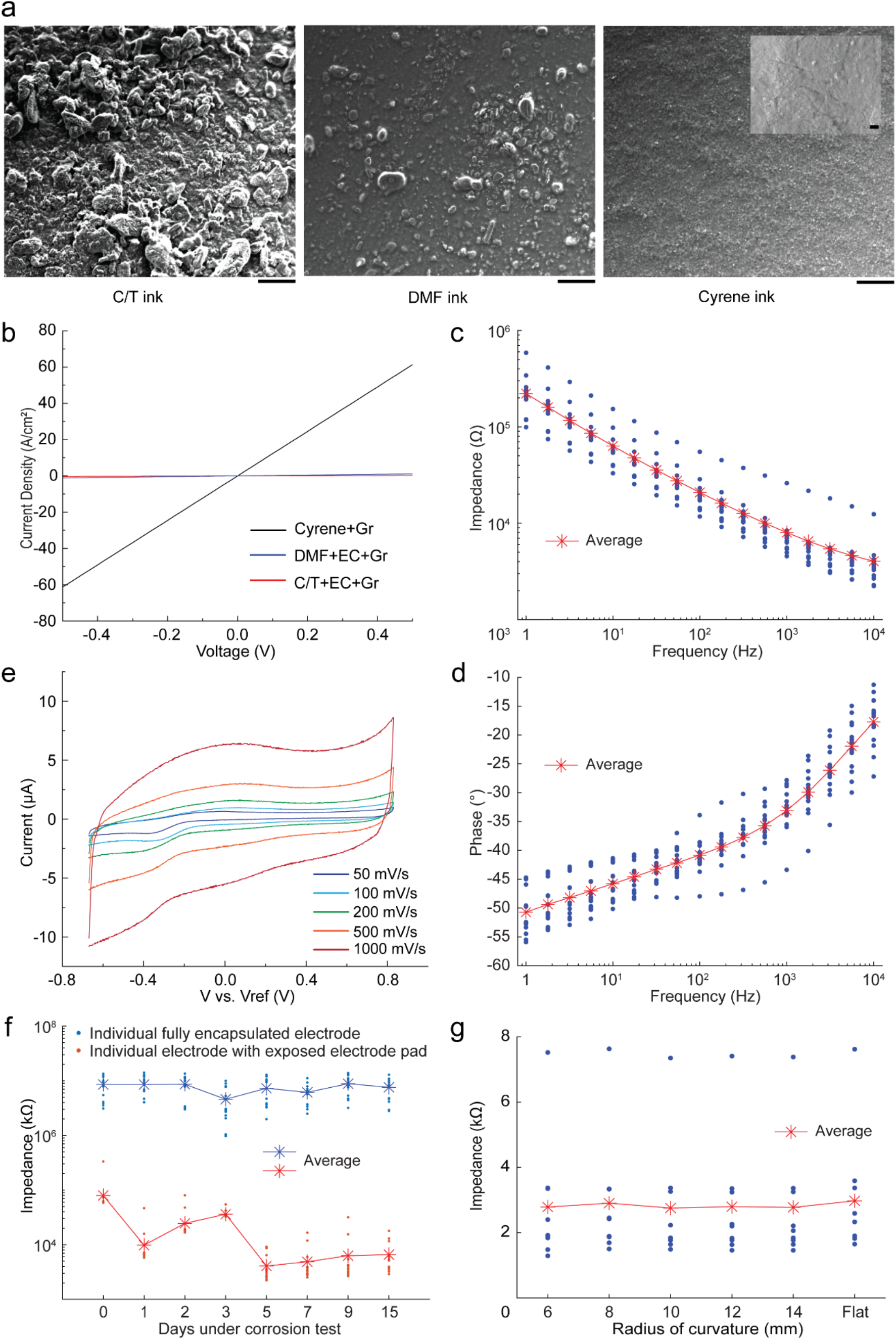
Material characterization of graphene films using three graphene inks and electrochemical characterization of graphene ECoG electrodes: **(a)** SEM images of the surface morphology of graphene films drop-cast onto a SiO2/Si substrate. The inset in the top right image shows the higher magnification image. Scale bars indicate 200 μm, while the scale bar in the inset indicates 2 μm. **(b)** Current density versus voltage characteristics of three graphene films. **(c)** Impedance magnitude at different frequencies ranging from 1 Hz to 10 kHz of electrodes made by stencil printing with Cyrene + graphene ink. **(d)** Phase angle at different frequencies ranging from 1 Hz to 10 kHz made by stencil printing with Cyrene + graphene ink. **(e)** Cyclic voltammetry of the graphene ECoG electrode in PBS at different scan rates in 50 to 500 mV·s^-1^. **(f)** Change in impedance magnitude as a function of time during an accelerated corrosion test. Each device has 16 electrodes with exposed electrode pads (red) and 16 electrodes with PDMS capsulated pads (blue). **(g)** The change of impedance magnitudes at 1 kHz as a function of radii of bending curvatures for nine identical straight graphene 10 mm long ECoG electrodes.

We next assessed the current density through the three graphene films at room temperature (**Fig. 2b**). At 0.5 V, the current density was measured to be ∼ 0.6 A/cm^2^ for the DMF+EC+Gr, ∼ 0.1 A/cm^2^ for C/T+EC+Gr, and ∼ 51 A/cm^2^ for Cyrene+Gr. This finding corresponds to the highest conductivity of the Cyrene ink sample (∼ 1.1 × 10^3^ S·cm^−1^). These results indicated that Cyrene was an adequate solvent for formulating the exfoliated graphene ink. The characteristics of small flake size and high conductivity of the exfoliated graphene in Cyrene indicate its potential for inkjet printing [38]. This work explored stencil printing to simplify the fabrication process for patterning the ECoG electrodes. Further experiments in this work involved devices made using graphene exfoliated in Cyrene.

### Benchtop characterization of stencil fabricated graphene ECoG electrode arrays

Cyclic voltammetry (CV) was used to evaluate the electrochemical stability of graphene electrodes [39]. The CV of the graphene electrode (n = 1) was measured at potentials with various scan rates ranging from 50 mV/s to 1000 mV/s. According to the shape of the plotted curves, both double-layer capacitance and pseudo-capacitance exist in the electrode (**Fig. 2e**) [40]. The capacitance value was extracted from the current (at 400 mV) vs. scan rate from the CV curve where the linear fit slope approximates the capacitance and was calculated to be ∼ 10 nF at 400 mV.

Electrochemical impedance spectroscopy (EIS) measured over a frequency range of 1 Hz to 10 kHz (**Fig. 2c**) was performed on a graphene ECoG device with 15 functional electrodes. The phase angle shows the typical capacitive behavior (near 80°) at 20 Hz but gradually becomes more resistive at higher frequencies (**Fig. 2d**). At high frequencies ranging from 1 kHz to 10 kHz, the relatively low impedance magnitudes are attributed to the stray capacitance and the Ohmic resistance of the electrolyte solution and interconnect [41]. The EIS of the stencil-printed graphene electrode presents similar impedance characteristics as a previously reported ECoG array of monocrystalline graphene layers [14].

An accelerated corrosion test was performed to evaluate the material stability of the graphene ECoG electrode (**Fig. 2f**). One graphene ECoG device with 16 exposed electrode pads, and a second device with 16 electrode pads fully capsulated by PDMS were submersed in 1X PBS maintained 60 °C, and the impedance was measured periodically. The electrodes in the fully capsulated device measured a consistently high impedance (> 1 MΩ) for 15 days, which indicates that the PDMS insulation layer is durable in the test. As for the device of exposed electrode pads, an initial reduction of the impedance was found in the first 3-4 days. After day 5, the average impedance was measured to be 3.97 ± 2.17 kΩ; after day 15, the average impedance was measured to be 6.49 ± 4.16 kΩ. These results indicate that the potential lifetime of the electrodes is at least 75 days for *in vivo* implantation [42].

The planar ECoG electrode array is typically bent to conform to the mouse’s convex dorsal cortex with an approximate radius of curvature of 12 mm [26]. Therefore, it is important to investigate the relationship between the electrode impedance and the bending radii of curvature. The impedances of 9 identical 10 mm long straight graphene electrodes were measured at 1000 Hz when the electrodes were placed on a flat surface and then bent at various radii of curvature using a custom measurement rig (**S. Fig. 3)**. No significant difference in the electrode impedance magnitudes on a flat surface or at different radii of curvature (14 mm to 6 mm) could be found (**Fig. 3g)**. Thus, the graphene electrodes patterned on the flexible PI substrates can be bent and conform to the dorsal cortical surface of the mouse brain with minimal effect on the electrode impedance.

**Figure 3.**
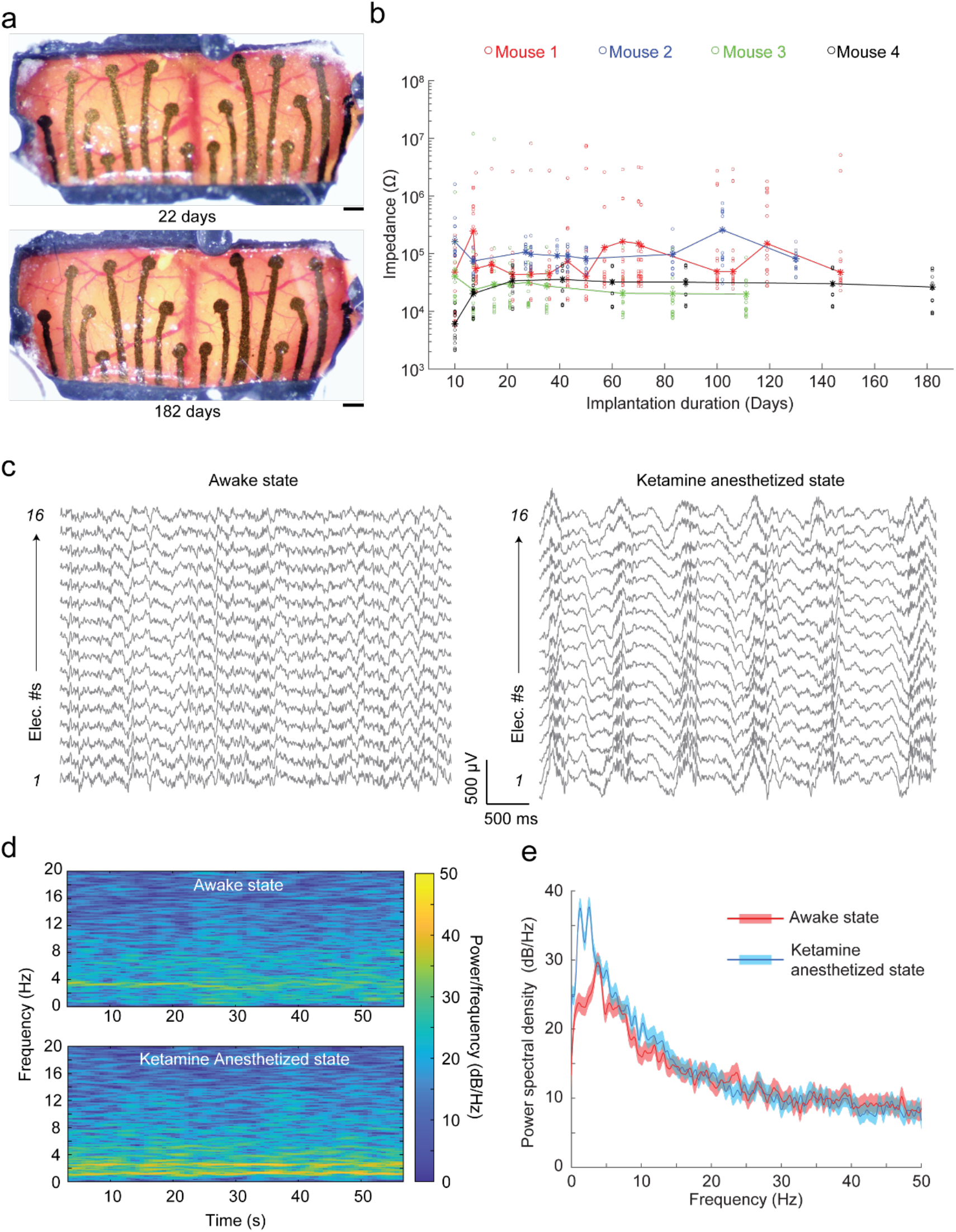
*In vivo* performance of the graphene ECoG devices: **(a)** Optical microscope images taken at 2 timepoints after chronic implantation of ECoG device on a C57BL/6 mouse (#4 shown in **(b)**. Scale bars indicate 500 µm. (**b)** Impedance magnitude at 1 kHz of 64 electrodes in 4 implanted mice. Dots indicate individual electrode impedance values. Stars indicate the average impedance of the 16 electrodes in each device. **(c)** Raw surface field potential recordings from the electrodes in awake and Ketamine anesthetized states. **(d)** Frequency spectrogram of electrode 5 shown in **(c)** in awake and anesthetized states. **(e)** Average power spectral density (PSD) of recordings of all electrodes shown in **(c)** in awake and anesthetized states.

### *In vivo* experiments

The graphene ECoG devices were implanted on multiple mice to evaluate the *in vivo* recording capabilities. The overall array dimension was ∼ 9 mm x 4 mm, covering most of the sensory cortex areas bilaterally (**Fig. 4b**). Representative microscope images of one such implanted mouse taken on days 22 and 182 after implantation are shown in **Fig. 3a**. No visible neuroimmune response such as Dural thickening and tissue encapsulation in these devices could be found. Most of the inner surface of the implant was covered by a PDMS insulation layer (**Fig. 1c**). PDMS itself is a biocompatible material for long-term cranial implants [43]. The longest duration of implantation assessed was 21 weeks. The impedances of the electrodes were periodically measured throughout the implantation (**Fig. 3b**). Across 4 mice, 61 electrodes (out of 64) remained functional, with an average impedance magnitude of 24.72 ± 95.23 kΩ (n = 61) at 1 kHz. Three electrodes lost their connections on mouse 1 at the measurement of day 10. ECoG electrodes were considered functional if impedance was lower than 1 MΩ [40], [41], [44]. The average electrode impedances of all 4 devices presented here remained less than 1 MΩ throughout implantation. Therefore, the graphene electrode arrays can act as robust electrical sensors for chronic study.

**Figure 4.**
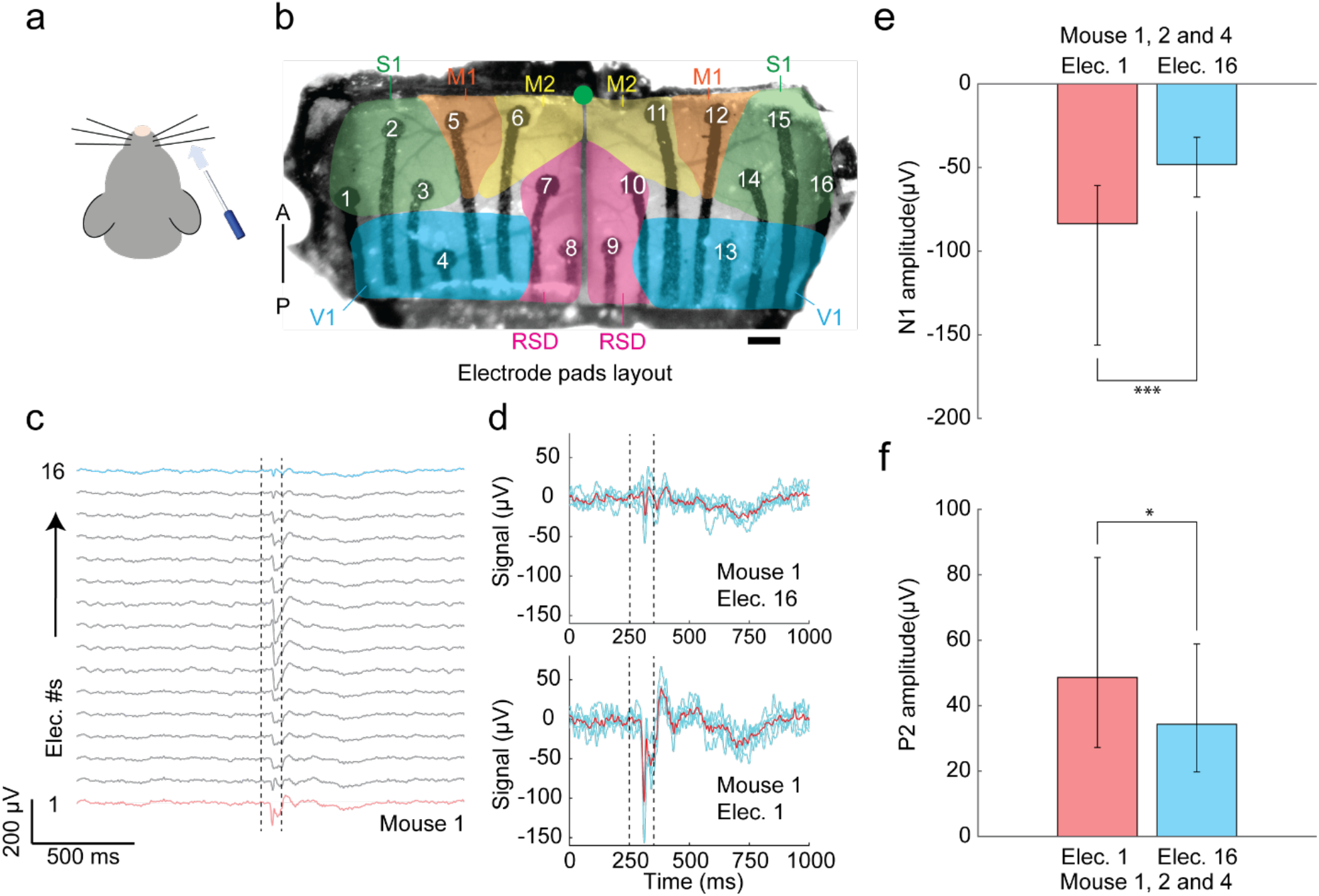
Sensory stimulus-evoked responses recorded by the graphene ECoG devices. **(a)** Air puff stimulus was applied to the right whiskers. **(b)** Locations of electrode pads: electrodes 1 - 3 and 14 - 16 were located in the primary sensory cortex (S1); electrodes 4 and 13 were located in the visual cortex (V1); electrodes 5, 6, 11, and 12 were located in the primary motor cortex (M1/M2). Electrodes 7, 8, 9, and 10 were located in the Retrosplenial cortex (RSD). Green dot indicates Bregma. Scale bar indicates 500 µm. **(c)** Average ECoG signals responses to air-puff stimuli (n = 138) in 20-min recording were collected from 16 electrodes of Mouse #1. Dotted lines indicate the on- and off-state of the air puff stimulus. Red line marks the signal of contralateral electrode 1, and blue line marks the signal of ipsilateral electrode 16 **(d)** Average ECoG signals captured by the contralateral electrode 1 (top) and ipsilateral electrode 16 (bottom) in response to repeated whisker stimulus (n = 5) in mouse 1. Blue lines indicate individual trials. Red line denotes the average signal response. **(e)** Comparison of surface field potential of depolarization’s peak amplitudes recorded by the contralateral electrode 1 vs. ipsilateral electrode 16 on mice 1, 2, and 4 (P < 0.001, *t-test*). **(f)** Comparison of local field potential repolarization’s peak amplitudes recorded by the contralateral electrode 1 vs. ipsilateral electrode 16 of the graphene ECoG on mice 1, 2, and 4 (P < 0.01, *t-test*).

ECoG recordings were performed in head-fixed mice while fully awake and under Ketamine-anesthesia to demonstrate the functional use of the graphene ECoG devices. The raw recordings obtained from all 16 electrodes in both states are shown in **Figure. 3c**. Consistent with previous observations [45]–[47], induction of the anesthetized state via Ketamine resulted in low-frequency oscillations at the delta frequency (0.5 – 4 Hz) throughout the cortex (**Fig. 3c**). These delta oscillations had signal peak-to-peak amplitudes of approximately 500 µV, much higher than the awake state’s amplitude of ∼ 200 µV (**Fig. 3c**). The spectrogram in **Figure. 3d** also presents a higher signal power density at 0.5 – 4 Hz at the anesthetized state, matching the power spectral density (PSD, **Fig. 3e**). The result indicates that all 16 channels of the device were functional to measure brain activity.

Furthermore, the graphene ECoG devices demonstrated the ECoG recordings of stimuli-evoked brain activity in response to sensory stimuli. The stimuli were the brief puffs of air (100 ms) given to the right whiskers of mice in a randomized fashion during the awake state under head-fixation (**Fig. 4a**). The stimuli resulted in whisking and the motor response observed in the experiments. Broad activation of most of the cortex in response to the stimuli was observed and possibly caused by the startle response (**Fig. 4c**). Channels 1 and 16 were individually located at the contralateral and ipsilateral sensory cortices (S1) (**Fig. 4b**). Channel 1 has higher signal amplitudes of negative peak N1 and positive peak P2 than channel 16 (**Fig. 4d**). Three animals (mouse #1, mouse #2, and mouse #4) were tested to study and analyze the signal difference between Channel 1 and 16. As shown in **Figure. 4e** and **f**, the average peak values N1 in electrodes 1 vs. 16 were - 83.60 ± 23.16 µV vs. - 48.20 ± 12.29 µV (p < 0.001, *t-test*) on all three mice; the average peak values P2 in electrode 1 vs. 16 were 48.64 ± 19.96 µV vs. 34.34 ± 11.28 µV (p < 0.05, *t-test*) on all three mice. Significantly stronger contralateral responses to the stimuli could be found in all three mice. Thus, the graphene ECoG electrode arrays could localize brain signals responding to stimuli.

## DISCUSSION

Here, we demonstrate for the first time a fully desktop fabricated, stencil-printed, liquid exfoliated, graphene-based, 16-channel ECoG electrode array on a flexible polyimide substrate, implanted on the rodent dorsal cortex for high-resolution neurophysiological recording. The highly stable graphene ink formulated by exfoliating graphene in a biocompatible solvent Cyrene [48], demonstrated a high conductivity ∼ 1.1 × 10^3^ S·cm^−1^. Alongside the electrochemical analysis, mechanical bending tests showed no significant change in electrode impedance at a flat surface and various radii of curvature ranging from 14 mm to 6 mm. Fully functional devices and the encapsulation of PDMS insulation were highly stable even under the accelerated aging environment, maintaining an average impedance of ∼ 6.49 ± 4.16 kΩ on day 15. Furthermore, the electrodes remained functional with an average impedance magnitude of 24.72 ± 95.23 kΩ (1000 Hz) throughout the implantation. Thus, our graphene ECoG arrays are robust neural interfaces that can be applied for chronic electrophysiology studies in mice.

To our knowledge, all the existing micrometer-scale ECoG devices are fully or partially fabricated using microfabrication or specialized techniques [42], [43], [49]–[56], resulting in high cost and low accessibility to neuroscience laboratories. In our method, the stencils determining the layout of the electrodes can be rapidly reconfigured using CAD tools and fabricated using desktop laser-cutters available in most university fabrication shops. The graphene ink can be formulated using standard lab equipment, and the patterned electrodes can be sintered on a laboratory hotplate. The insulation layer can be deposited by a homebuilt, microcontroller-based spinner or a low-cost off-the-shelf spin coater. We have created the first fully desktop fabricated, flexible micrometer-scale ECoG array that can be chronically implanted in mice. These results point a way forward for creating robust, open-source, flexible neural interfaces that can be widely used in basic and translational neuroscience research.

We identified some limitations to the proposed approach. First, the minimum feature size is limited by the resolution of the desktop laser cutter to create the stencils. The ECoG devices with an inter-electrode (center to center) distance of ∼ 500 µm can be patterned using the laser cutter for a stable fabrication quality, limiting the overall number and density of electrodes incorporated within a device. Second, the graphene ink needs to be annealed at high temperatures (> 300°C), limiting the substrates used for supporting the graphene electrodes. As an alternative, photonic sintering can be used instead of thermal annealing, allowing the use of other materials as substrates. For instance, the electrodes could be patterned on transparent polymers such as polyethylene terephthalate (PET) to create devices for simultaneous imaging and ECoG recording [11]. Though transparent graphene ECoG has been developed, it still requires cleanroom equipment [49]. Our desktop-fabricated stencil printing method does not apply to building transparent graphene ECoG, but off-shelf transparent conductive PEDOT:PSS ink can be a potential alternative [8] for making fully transparent ECoG.

Several directions can be pursued in the future, building upon the graphene ECoG devices presented in this work. The size of the implant can be extended to much larger regions of the brain [26], [27] so that the whole visual cortex and motor cortex can be included for performing ECoG recording over most of the cortical surface of the mouse brain [11]. The low and stable impedance of the electrodes can be leveraged for precise cortical micro-stimulation [14]. Miniaturization of electronic interface circuits can also allow deployment in freely behaving animals, potentially combined with imaging instrumentation for simultaneously tracking large-scale calcium dynamics [27]. Further, graphene has shown utility in passive sensing of electrical potentials and functionalizing to highly sensitive biochemical sensing [57]. Overall, our method can be scaled to mass manufacturing of flexible graphene-based biosensors at a low cost with a myriad of applications in biological sensing, such as electrocardiography [58], electromyography [59], and peripheral nerve interfacing [60].

## METHODS

### Graphene inks formulation

The graphene inks were formulated by bath ultra-sonication of graphene powder, which was ground using a mortar pestle from a graphite rod (496561, Sigma Aldrich), in a mixture of Cyclohexanone:Terpineol (C/T) (398241 and 814759, Sigma Aldrich) at a ratio of 7:3, DMF(319937, Sigma Aldrich), and Cyrene (807796, Sigma Aldrich). An initial concentration of Ethyl Cellulose (EC) (EC 200646, Sigma Aldrich) at 2.5 wt% and 10 mg/mL graphene powder were then added in 10 mL C/T and DMF, treated for 2 hours in the Branson Bath Sonicator (CPX2800, Fisher Scientific) at 30 °C. No EC was added to the Cyrene ink as it produces excess bubbles resulting in bad drop casting. All three dispersions were kept idle for 24 hours to precipitate the larger particles, which left pale gray precipitation at the bottom of the vials. The supernatant (∼ 8 mL) inks were extracted and stored in clean vials.

### Graphene sample preparation for Electrical Measurement

To carry out the initial electrical characterization of the graphene inks, high integrity metal contacts composed of silver (*Ag*) were first patterned with photolithography and deposited with an e-beam evaporator on the PI film, followed by drop-casting of 0.5 mL graphene ink in a rectangular area of 10.5 mm x 2 mm using a stencil (**S.Fig. 4**). The sample was annealed at 350 °C for 90 minutes. A probe station with parameter analyzer (B1500A, Agilent Technologies) was used to conduct 2-point-probe voltage-controlled measurements and data extraction.

### Electrode array design and assembly

#### Electrode array design

The layout of the graphene electrode array was rendered in computer-aided design software (SolidWorks 2021, Dassault) (**Fig. 1b**). The center-to-center distances between 2 neighboring electrode pads ranged from ∼ 600 µm to 1 mm. The overall layout of the electrode array was designed to cover an area of ∼ 9 mm long, extending bilaterally, with a width of ∼ 4 mm posterior to Bregma (**Fig. 5b**). Electrodes 1-3 and 14-16 were in the primary sensory cortex (S1); electrodes 4 and 13 were located in the visual cortex (V1); electrodes 5-6 and 11-12 were located in the primary motor cortex (M1/M2), and 7-10 were located in Retrosplenial cortex (RSC).

#### Laser-cut stencils

To fabricate the stencils, a piece of 50 µm-thick insulating polyimide (PI) tape (High Temperature Tape, Bertech) with a size of 25.4 mm by 160 mm was adhered to a stainless-steel sheet and subsequently patterned by a CO_2_ laser cutter (PLS6.140D, Universal System) (**Fig. 1a**). We tested the laser cutter’s cutting precision on the PI tape and found that the inter-electrode pitch (center to center distance) of ∼ 500 µm with power settings of 10% and speed settings of 100% can achieve the stable stencil (**S.Fig. 1b**). The electrode via stencil was transferred to a polyimide film substrate using a scotch tape transfer method modified from previous work [61].

#### Graphene ink deposition

The graphene ink was manually drop-cast on the laser-cut mask and annealed at a high temperature (∼ 350 °C) for 90 minutes. To evaporate the solvent completely, the hotplate temperature was ramped at ∼ 10 °C/minute from 100 °C to 330 °C to evaporate the solvent completely. After annealing, the mask was carefully removed, leaving the patterned graphene channels on the polyimide film. A similar procedure was used to apply a sacrificial layer of Pluronic (F-127, Sigma Aldrich) after manually aligning the electrode pad stencil to the graphene electrodes. The sample was kept overnight at room temperature to solidify the Pluronic. Diluted Polydimethylsiloxane (PDMS) was spin-coated at 1500 rpm for 15s using a home-built desktop spin-coater (**S.Fig. 5**), then cured at 30 °C temperature for 15 minutes. For making the diluted PDMS, the tert-Butanol (471712, Sigma) was warmed at 45 °C and mixed with PDMS and SYLGARD 184 curing agent (761036, Sigma) at the weight ratio of 50:10:1. The Pluronic was released with a gentle flow of hot water at 45 - 60 °C to expose the electrode pads. Then the sample was annealed again at 100 °C for 30 minutes.

#### Implant assembly

The graphene ECoG device is an assembly of four major components: the stencil fabricated ECoG array, a 3D printed cranial window frame, a flexible printed circuit board (PCB) connector, and a 3D printed reinforcement clamp. The cranial window frame and the reinforcement clamps were 3D-printed using a stereolithography 3D printer (Form 2, Formlabs). The PCB connector was custom-designed in Eagle (Autodesk Inc.) and fabricated by an online PBC manufacturer (PCBWay.com). To assemble the device, the ECoG electrode array was first bonded to the gold soldering pads on the flexible PCB connector using a conductive adhesive (8331 Silver Epoxy Adhesive, MG Chemical) (**S. Fig. 2**) by applying it manually with a sharp object. The interface was further mechanically reinforced using the 3D-printed clamp and followed by encapsulation with clear epoxy adhesive (Scotch-Weld Epoxy Adhesive DP100 Plus, 3M). The recording area of the stencil fabricated graphene electrode arrays was bonded to a 3D printed cranial window frame using epoxy adhesive (DP1000 Plus Clear, 3M) adapted from our previous work [26], [27] to realize the device illustrated in **Fig. 1f**. The cranial window defined a total recording area of ∼ 4 mm x 9 mm, with a radius of curvature of 10 mm, to allow conformal implantation over the dorsal cortex immediately posterior to Bregma (**Fig. 5b**). A fully assembled device has a mass of ∼ 1.3 g.

### Benchtop testing of fully assembled devices

CV, EIS, and accelerated corrosion tests were performed on fully assembled graphene ECoG devices. EIS and CV measurements were conducted using a potentiostat (1010E, Gamry Instruments), with an Ag/AgCl reference electrode (Gamry Instruments) and a platinum wire as a counter electrode (1.0 mm diameter, Premion, 99.997%, Alfa Aesar) in room temperature 1X Phosphate Buffered Saline solution (D1408 PBS 10X, Sigma-Aldrich). EIS was performed with a 50 mV excitation voltage from 10 kHz to 1 Hz. Ten measurements were taken per decade. The CV was performed at varying scan rates from 50 – 1000 mV/s, with voltage limits set at ± 0.75 V vs. the open circuit potential.

The accelerated corrosion test was performed [62] by immersing 2 ECoG devices (one device has 16 exposed electrode pads, and one device has 16 fully PDMS-capsulated electrode pads) into 1X Phosphate-buffered saline (D1408 PBS 10X, Sigma-Aldrich) at an elevated temperature of 60 °C which is equivalent to a five-fold accelerated corrosion process at body temperature of 37 °C [42]. Impedances of the device were measured using the interface board (RHD2000, Intan Technologies) at 1000 Hz daily for 2 weeks.

Bending testing was conducted by creating 9 straight, parallel graphene electrodes using the same ECoG stencil printing technique. To mimic various degrees of bending after implantation, the electrodes were placed on a flat surface and custom-built acrylic structures (**S.Fig. 3**) with radii of curvature of 6 mm, 8 mm, 10 mm, 12 mm, and 14 mm, when their impedance was measured using the interface board (RHD2000, Intan Technologies Inc.) at 1000 Hz.

### Surgical implantation

All animal experiments were approved by the University of Minnesota Intuitional Animal Care and Use Committee (IACUC). 9 C57BL/6 mice (5 males and 4 females) at the age of 12-30 weeks were used in this study. Initially, 5 mice were implanted with the graphene ECoG devices during the optimization of the electrode layout and overall device design. Mice were housed in a 14hr light/10hr dark cycle in rooms maintained at 20-23 °C and 30-70% relative humidity. Mice had ad libitum access to food and water. Mice were given preemptive doses of 2mg/kg of sustained-release buprenorphine (Buprenorphine SR-LAB, ZooPharm) and 2 mg/kg of meloxicam for analgesia and preventing brain inflammation respectively. Mice were anesthetized 30-60 minutes after the initial analgesia dosage using 1-5% isoflurane anesthesia in Oxygen. Eye ointment (Puralube, Dechra Veterinary Products) was applied to the eyes. The scalp was shaved and cleaned. Once the mice were fixed in a stereotax (900LS, Kopf), the scalp was sterilized by repeatedly scrubbing Betadine and 70% Ethanol solution (3 times). Next, the scalp was removed using surgical scissors. The tissue and fat under the scalp were subsequently cleared using a micro curette (# 10080-05; Fine Science Tools). Partial temporalis muscle wrapping around the skull was carefully removed using a scalpel to expose the squamosal area. After drilling a ∼ 300 µm-diameter hole on the squamosal suture, a stainless-steel bone screw (#FF000CE094, JI MORRIS Company) tied to a 26-gauge stainless-steel reference wire was tightened in the hole firmly. A large craniotomy was immediately performed using a high-speed dental drill following a rectangular path approximately 4 mm x 9 mm posterior to Bregma. After the drilling, the skull piece was removed from the dorsal cortex using two micro curettes holding its lateral edges. A gauze pad soaked in sterile saline was gently placed on the exposed brain to keep it moist.

The graphene ECoG device was sterilized by immersion in 70% Ethanol for 2 min and subsequently rinsed thoroughly with sterile saline. The periphery of the craniotomy was cleaned using a pointed cotton tip after removing the gauze pad. The window frame was gently placed on the exposed brain. Surgical adhesive (Vetbond, 3M) was applied around the edges of the window frame to bond the window frame to the skull. After the adhesive was cured, a customized waterjet-cut Titanium headplate was fastened on the implant with a #0-80 screw. The implant was cemented to the skull using opaque dental cement (Metabond, Parkell Inc.). Mice were transferred from the stereotax to a heated pad (catalog no. 72-0492; Harvard Apparatus) for recovery from the anesthesia and were transferred to a clean cage partially located on a warming pad once they were fully ambulatory. 2 mg/kg meloxicam was administrated twice per 24 hr for 72 hr post-surgery. Mice recovered for 7 days before any *in vivo* experiment.

### *In vivo* electrophysiology

#### *In vivo* electrode performance test

The impedances of electrodes in the implanted devices were measured periodically (up to 182 days). Animals were transferred from their home cage and affixed under anesthesia in a custom head-fixation device adapted from previous studies [26], [63]. Subsequently, the FPC connector on the amplifier was connected to the PCB connector of the ECoG device. Impedance measurements were acquired at 1000 Hz.

#### Awake and anesthetized spontaneous recordings

Mice were head-fixed on the treadmill, and 4 minutes of spontaneous recordings were acquired when the animals were fully awake. All the signals were recorded through the RHD 2000 interface board at a sampling rate of 20 kHz. Mice were administrated a cocktail of Ketamine (100 mg/kg) and xylazine (10 mg/kg) while head-fixed. A heated pad was placed under the animal to maintain body temperature, and mice were supplemented with Oxygen. Once the animal was fully unresponsive to toe pinch stimulus, multiple 4-minute-long spontaneous recordings were acquired. Once fully recovered from anesthesia, mice were removed from the head fixation apparatus and transferred back to their home cage.

#### ECoG recordings in response to sensory stimuli

Brief air-puff stimuli to the awake animal’s right whiskers were applied through a blunt 24-gauge needle to measure ECoG responses to sensory stimuli. The air-puff stimuli were generated using a compressed air source. The duration of the stimulus (∼100 ms) was controlled using a microcontroller (Arduino Uno, Adafruit) actuated solenoid valve. Airpuff stimuli were separated by 8 to 10s, and 24-30 stimuli were given to each mouse.

### Data analysis

All the data, including electrode impedances and ECoG signals, were collected and converted using the interface board’s software (RHD2000 interface, Intan Technologies Inc.). The raw ECoG signals were first down-sampled to a moving average of 2 kHz and low-pass filtered using an elliptic filter with a passing band of 100 Hz. Custom scripts in MATLAB (MATLAB 2020b, Mathworks) were used to analyze and plot the data.

## AUTHOR CONTRIBUTIONS

SBK conceived the overall project. JH, RFH, AT, and SBK conceptualized the devices. JH developed the electrode array designs, laser-cut stencil technique, implant, and electrode interface. JH also designed and performed the *in vivo* experiments, surgical implantation, and data analysis with the help of ZN. RFH developed the recipe for the graphene inks and characterized the properties. RFH performed the electrical & material characterization and analysis. RFH also engineered and developed the stencil fabrication process of the ECoG electrode array and cleanroom fabrication of the test samples. JH and RFH performed the electrode bending test. PDD carried out CV and EIS experiments with the help of RFH and JH. ML designed and built the spin coater. JH, RFH, PDD, and SBK wrote the paper. All authors revised and contributed to the final manuscript.

## ACKNOWLEDGEMENTS

SBK and SLS acknowledge NINDS Award #R0NS111028. SBK acknowledges Brain Initiative Award R42NS110165. Parts of this work were carried out in the Characterization Facility, at the University of Minnesota, which receives partial support from the NSF through the MRSEC (Award Number DMR-2011401) and the NNCI (Award Number ECCS-2025124) programs. We acknowledge Javier Garcia Barriocanal for assistance with the XRD measurements. PDD was supported by NSF IGERT Award DGE-1069104. Skylar Fausner for assistance with animal preparation. We thank Skylar Fausner and Beatrice Gulner for useful comments and critiques of the paper.

## INSTITUTIONAL APPROVAL

All animal experiments described in this paper were approved by the University of Minnesota’s Institutional Animal Care and Use Committee (IACUC).

## Supplementary Figures

**S. figure 1.**
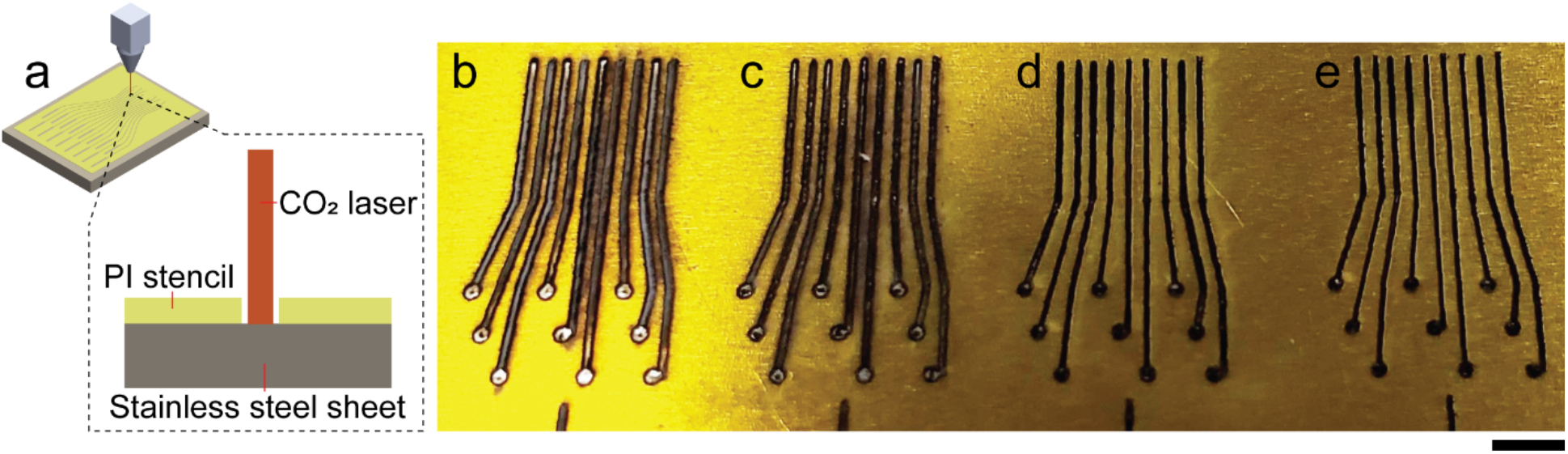
Laser-cut stencil characterization. **(a)** The 2D schematic of laser cutting the stencil of electrode vias. The stencil was cut using a commercial CO2 laser cutter (PLS6.140D, Universal System) through the PI tape supported by a stainless-steel sheet. **(b)-(e)** Images of laser cut stencils at different power and speed **(b)** A laser power setting of 10% and speed setting of 50% failed to create a distinct boundary between 2 neighboring electrodes. **(c)** A laser power setting of 10% and a speed setting of 100% created fully etched vias with distinct boundary between electrode tracks. **(d)** At a laser power setting of 5% maximum and speed setting of 100% maximum, the laser could fully cut through the PI tape but had carbonized residue filling in the stencil, which was hard to remove. **(e)** At a laser power setting of 2.5% maximum and speed setting of 100% maximum, the laser could not fully cut through the PI tape. The scale bar indicates 2 mm. We chose the settings in (c) for further experiments.

### Laser-cut stencil optimization

to achieve the most stable design of the electrode array, the laser cutting power and speed determine whether the adjacent electrodes are isolated from each other without any interconnects. **S. Figure 1a** presents the animated schematic, in which the CO_2_ laser cut through the PI stencil supported by a stainless-steel sheet. The optimized power at 10% and speed at 100% experimentally achieve the best stencil of the electrode’s center to center distance of 500 µm (**S.Fig 1b – e**).

**S. figure 2.**
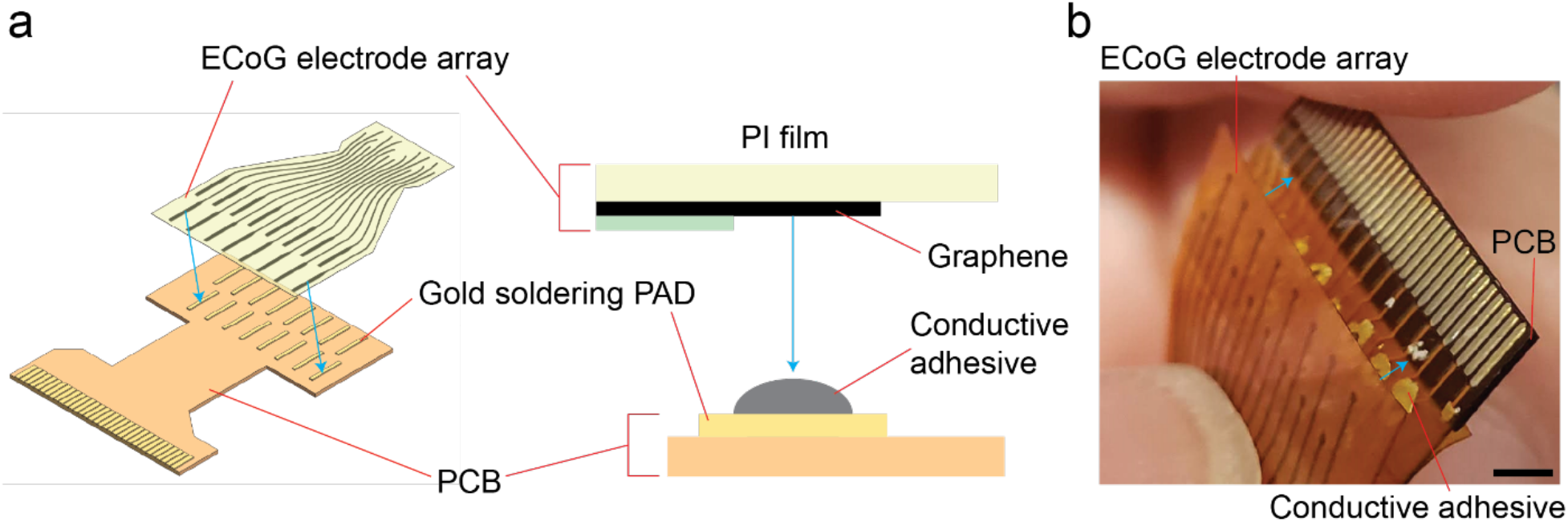
bonding between PCB and ECoG electrode array. **(a)** 2D and 3D schematic of the bonding between the graphene ECoG array and interface PCB using conductive adhesive. **(b)** Photograph illustrating graphene ECoG array and interface PCB at the bonding sites. The scale bar indicates 2 mm.

### Bonding between ECoG electrode array and PCB interface

The PCB’s metal pads are normally used for soldering. Silver epoxy (8331 Silver Epoxy Adhesive, MG Chemical) was applied individually on each metal pad and subsequently bonded to the graphene electrode (**S.Fig. 2a**,**b**).

**S. figure 3.**
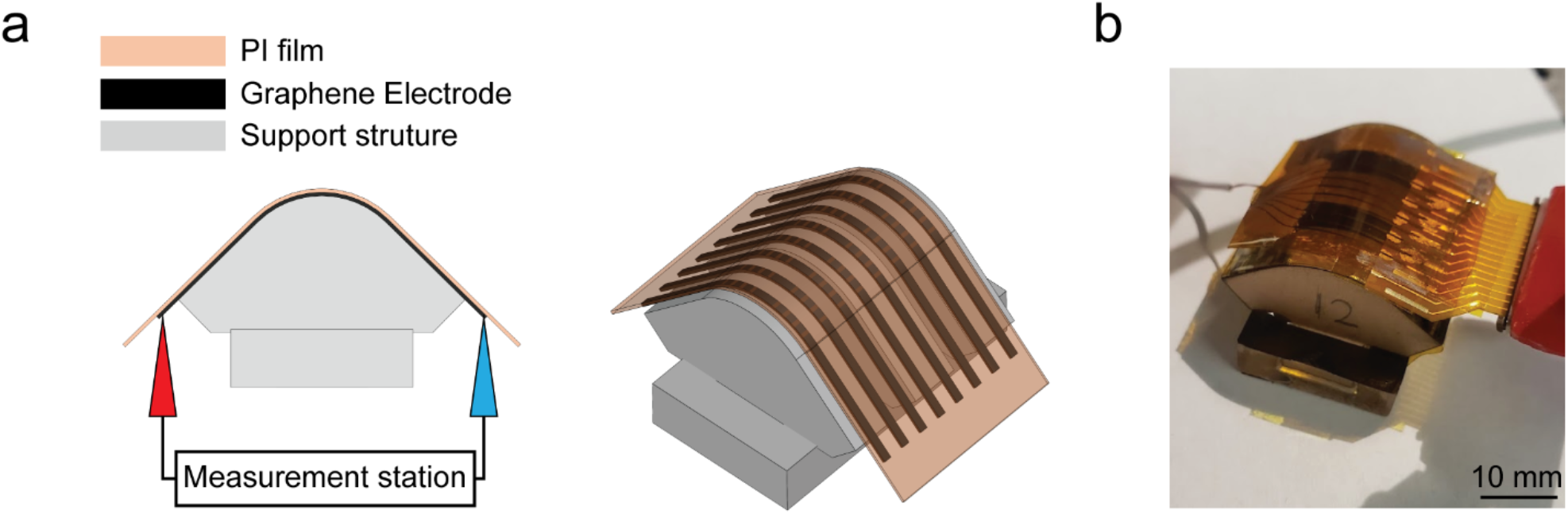
Bending test. **(a)** The 3D and 2D schematic of the bending test setup. 3 identical electrodes built by the same fabrication procedure of the graphene ECoG electrode array were conformed to the custom supporting structure with a curvature. The electrode length is 15.7 mm. The electrode width is 100 µm. **(b)** Photograph of the measurement setup.

### Bending test

to evaluate the electrode performance under static bending stress, 9 straight electrodes built using the same ECoG stencil printing technique were placed on the support structures with different radii of curvature. The impedance was measured using the interface board (RHD2000, Intan Technologies Inc.) (**S.Fig 3a, b**).

**S. figure 4.**
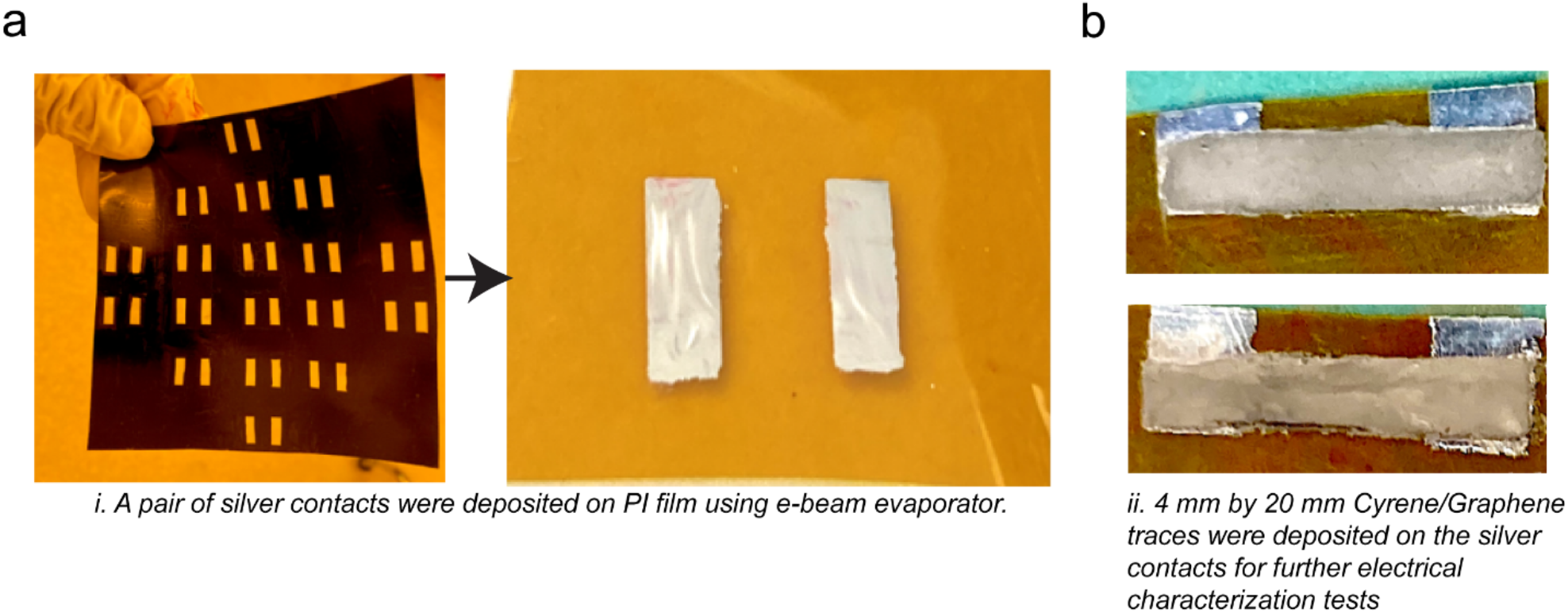
Sample preparation for electrical characterization: **(a)** A pair of silver contacts were deposited on PI film using e-beam evaporator through a lithography mask. **(b)** 4 mm x 20 mm graphene trace was patterned on the silver contact for electrical characterization tests.

### Sample preparation for electrical characterization

To fabricate the *Ag* contacts on the PI film, a dark mask was first printed on a transparent PET (Polyethylene terephthalate) substrate (**S.Fig 4. a**) using a regular desktop printer. The AZ5214 photoresist was spin-coated at 4000 rpm for 30 s on the PI film and baked at 110°C for 90 s on a hot plate. It was then exposed to the dark mask with UV light using a mask aligner for 30 s. It was then developed in the AZ developer and after developing, a clear transparent pattern on the PI film was visible. Then the e-beam evaporator was used to deposit the *Ag*, which deposited a uniform coating, which later went through a lift-off process with acetone solvent to achieve the desired pattern of *Ag* contact. ∼ 0.5 mL of three graphene inks were drop cast and annealed at 350°C for 90 minutes to create the samples.

**S. figure 5.**
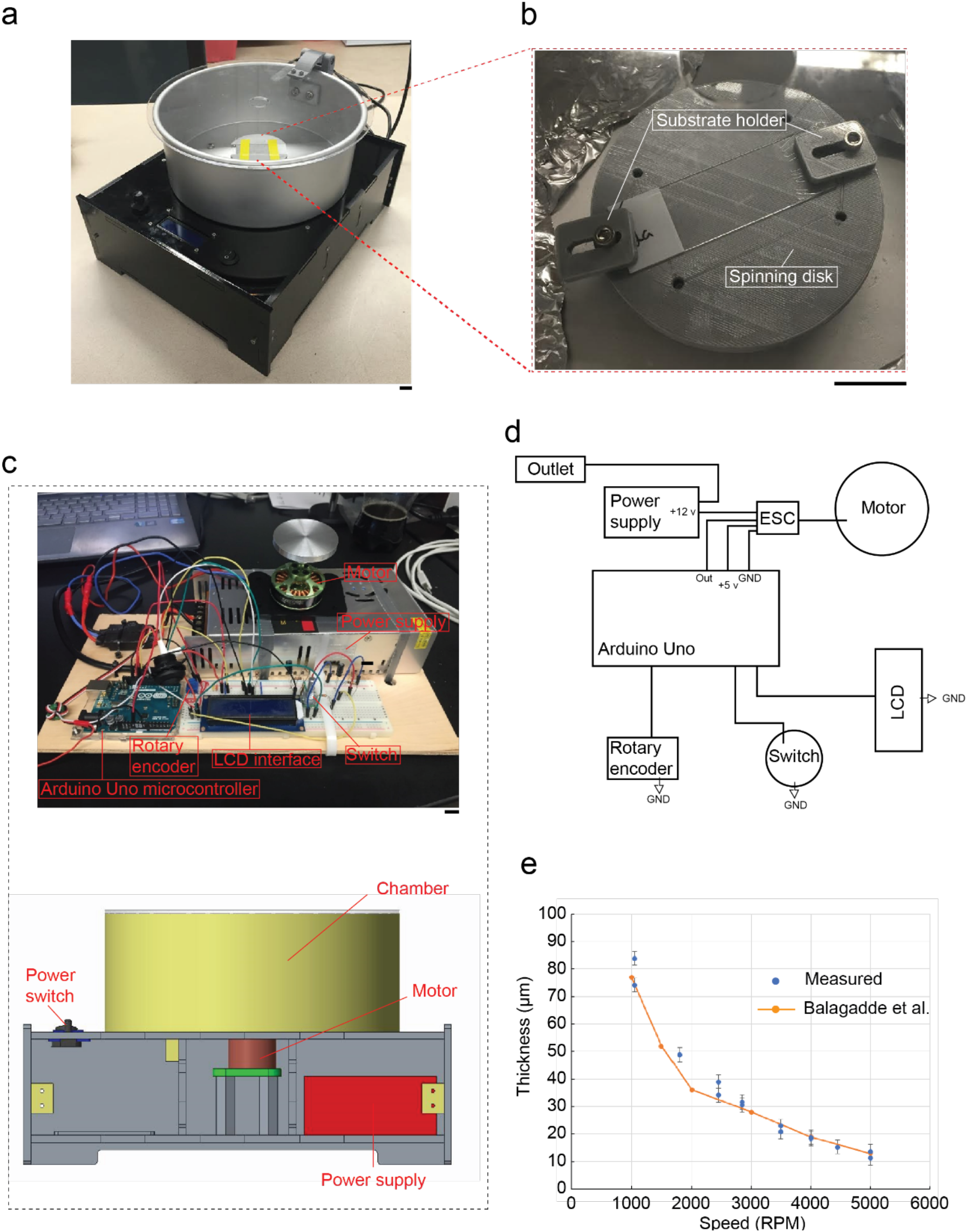
Custom-built spin coater: **(a)** Photograph of a custom-built microprocessor-controlled spin coater. **(b)** The spin coater has one spinning disk with 2 substrate holder clamps that hold the substrate for spin coating. **(c)** The major components of the spin coater include a microcontroller, rotary encoder, liquid crystal display (LCD) interface, power switch, power supply, motor, and chamber. **(d)** Circuit layout. ESC in the schematic abbreviates electronic speed controller. **(e)** The relationship between spinning speed and PDMS thickness was obtained using the custom-built spin-coater compared to results obtained using a commercial spinner [1].

### Custom-built spin coater

the spin coater was designed to be powered by a 12-volt power supply and run with an Arduino microcontroller (**S.Fig 5. d**). The use of an Arduino is convenient due to its ability to be easily programmed and tested. The motor can operate in a specific rpm window of about 2000 to 8000 RPMs; an electronic speed controller (ESC) can supply a pulse-width modulation signal from the Arduino microcontroller to control the motor (**S.Fig. 5d**). The device is mainly controlled by a rotary encoder (**S.Fig. 5c**). A standard 16×2 LCD was used to display adjustable options and a timer and RPM counter when the spin-coater was working (**S.Fig. 5c**). To allow easy cleaning, acrylic sheets for the enclosure and a removable cake pan were selected as the spinning chamber (**S.Fig. 5a**). The sample can be placed on the spinning disk and clamped by a pair of substrate holders (**S.Fig. 5b**)

**S. figure 6.**
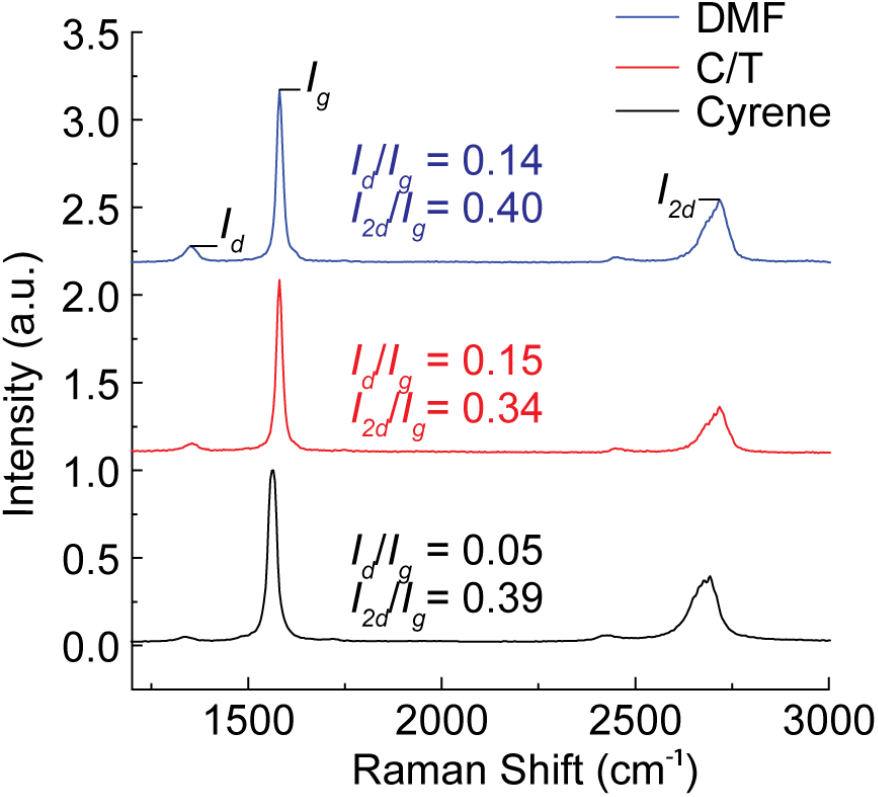
Raman spectrum: Raman spectrum of the three graphene films at room temperature, showing distinct Raman D, G, and 2D-band peaks, where the Id/Ig ratio was lower for the film originated formed from Cyrene Cyrene-exfoliated graphene ink.

### Graphene sample preparation for Raman Spectroscopy

0.5 mL of the prepared graphene ink was extracted from the vial using a disposable pipette and released onto a Si/SiO_2_ substrate, followed by spin coating at 1000 rpm for 20 sec using a custom-built microcontroller-based spinner (**S.Fig. 5**). To expose single/multilayer graphene flakes and minimize stack-up of multiple flakes, the sample was then annealed at 350 °C for 90 minutes. A 532 nm wavelength laser was used to perform Raman Spectroscopy measurements. The laser power was set to an initial setting of 0.10 mW and increased gradually to 16 mW. The measurement acquisition time was fixed at 0.1 s to minimize any local heating that could be caused by the laser [1].

**S. figure 7.**
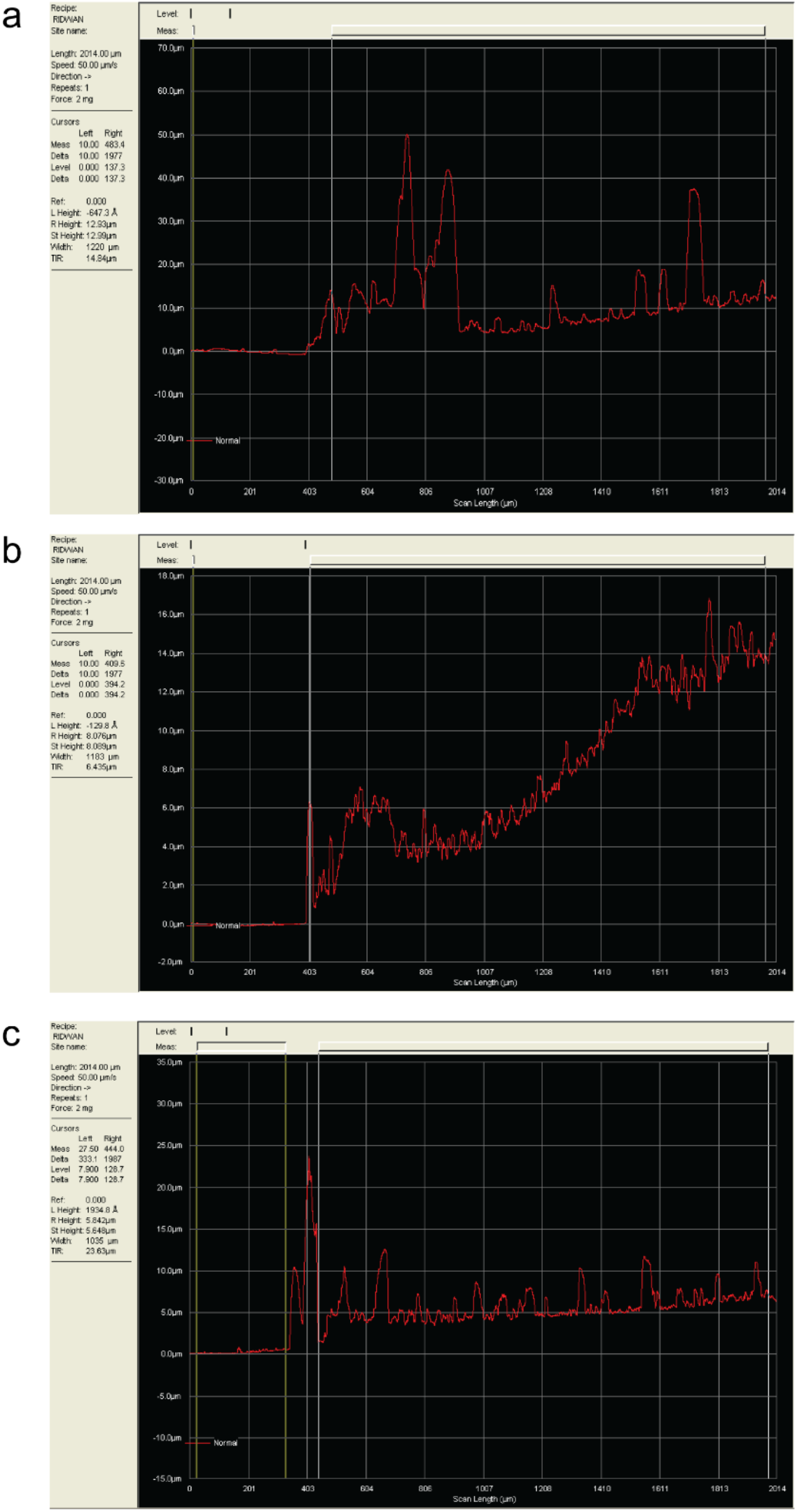
Graphene electrode surface profiling: **(a)** C/T+EC+Gr electrode. **(d)** DMF+EC+Gr electrode. **(c)** Cyrene+Gr electrode.

We used a surface profiler (K1-P16, KLA-Tencor) to measure the Graphene electrode surface profile.

